# Lake Erie fish safe to eat yet afflicted by algal hepatotoxins

**DOI:** 10.1101/2022.06.07.495188

**Authors:** René S. Shahmohamadloo, Satyendra P. Bhavsar, Xavier Ortiz Almirall, Stephen A. C. Marklevitz, Seth M. Rudman, Paul K. Sibley

**Affiliations:** School of Biological Sciences, Washington State University, 14204 NE Salmon Creek Ave, Vancouver, WA, 98686, United States; School of Environmental Sciences, University of Guelph, 50 Stone Rd E, Guelph, ON, N1G 2W1, Canada; Environmental Monitoring and Reporting Branch, Ontario Ministry of the Environment, Conservation and Parks, 125 Resources Rd, Toronto, ON, M9P 3V6, Canada; Department of Physical & Environmental Sciences, University of Toronto, 1065 Military Trail, Toronto, ON, M1C 1A4, Canada; IQS School of Engineering, Universitat Ramon Llull, Via Augusta, 390, 08017, Barcelona; Lake Erie Management Unit, Ontario Ministry of Northern Development, Mining, Natural Resources and Forestry, 320 Milo Road, Wheatley, ON, N0P 2P0, Canada

**Author notes:** Address correspondence to: René S. Shahmohamadloo, PhD, School of Biological Sciences | Washington State University, 14204 NE Salmon Creek Ave | Vancouver | Washington | 98686 | United States. **Author Contributions:** P.K.S., and S.P.B. designed the research. S.A.C.M. conducted field sampling. R.S.S., S.M.R., and X.O. analyzed the data. R.S.S. and S.M.R. wrote the paper. All authors read, amended, and approved the final manuscript. **Competing Interest Statement:** The authors declare no competing interests.

**Keywords:** Fish, Harmful Algal Bloom, Human Health, Wildlife Management, Great Lakes

## Abstract

Microcystin toxins from harmful algal blooms (HABs) can accumulate and persist in fish, raising dual concerns about human health risks from consumption and the potential for detrimental impacts on fish populations. However, there are fundamental unknowns about the relationship between HABs and fish populations driven by a lack of field information on toxin accumulation and retention over space and time. We conducted a field study to assess human health risks from consuming fish caught across all life stages of a HAB and to determine the pervasiveness of potentially harmful levels of microcystins on fish populations. We collected 190 fish in 2015 and 2017 from Lake Erie, a large freshwater ecosystem that is highly productive for fisheries and is an epicenter of HABs and microcystin toxicity events. Muscles and livers were analyzed for total microcystins, which was used to conduct a human health risk assessment for comparison against fish consumption advisory benchmarks available for Lake Erie. We find low human health risk from muscle consumption following the World Health Organization’s safety thresholds. However, all fish across capture dates had microcystins in their livers at levels shown to cause adverse effects, suggesting a pervasive and underappreciated toxic stressor. These data demonstrate that microcystins are retained in fish livers well beyond the cessation of HABs and calls for additional research to better understand the effects of sublethal toxic exposures for fish population dynamics, conservation, and related ecosystem services.

## Introduction

It has been over 50 years since the United States and Canada committed to restoring and protecting the waters of the Great Lakes (1). Yet, harmful algal blooms (HABs) continue to pose a tremendous risk to freshwater biodiversity conservation (2), precipitating a concerted effort to explore opportunities for intervention to mitigate the emerging threats to populations, communities, and ecosystems (3). The frequency and severity of HABs are increasing due, in part, to global changes in climate and anthropogenic eutrophication (4). The economic losses from HABs are estimated to exceed US$4.2 billion annually in the United States and Canada combined, due to deleterious effects on essential goods and services, including drinking water, recreation and tourism, and fisheries (5, 6). Food recalls linked to tissue accumulation of HAB toxins in fish and poisoning cases in humans and animals are also rising (7). Among the most widespread of these toxins are microcystins, a class of hepatotoxins that can be tremendously detrimental to human health (4, 8) and have been measured in fish tissues at levels exceeding the World Health Organization’s (WHO) tolerable daily intake (TDI) value (0.04 μg kg^-1^ bodyweight per day) (9). However, the true concentration of microcystins in fish tissues is debated because existing work has relied heavily on the enzyme linked immunosorbent assay (ELISA), an inaccurate method which was not designed for fish tissues and is prone to false positives (10, 11). Consequently, the actual risks to human health from consuming fish that encounter and accumulate microcystin-HABs remain poorly characterized.

Beyond concerns for human health, microcystins can adversely impact key aspects of fish life history including growth and recruitment (12). The overall literature on the magnitude of the effects of microcystins on fish health is unresolved; some studies suggest fish can effectively eliminate microcystins (13), while others suggest they cannot (14). However, recent mechanistic work has demonstrated that fish can exhibit stress responses even after being removed from a toxic bloom (15). There is also emerging pressure to ensure HABs do not deter the development of aquaculture and fisheries to feed an expanding human population (7, 16). Fish species inhabiting Lake Erie, a shallow freshwater ecosystem known for consistent and toxic HABs (17, 18), are particularly at risk. Determining the extent of microcystin accumulation and retention across time and species is therefore a crucial step towards understanding the impacts of HABs on the health of fish and fisheries.

We analyzed 190 Walleye (*Sander vitreus*), White Bass (*Morone chrysops*), White Perch (*Morone americana*), and Yellow Perch (*Perca flavescens*) from Lake Erie in 2015 and 2017 pre-, during, and post-HAB periods occurring in the western, west central, and east central basins. We dissected and analyzed the total microcystin content of: 1) muscles, which is of socioeconomic and cultural significance to Great Lakes fisheries and communities (6), and 2) livers, which is the primary target organ of microcystin toxicity and driver of fish mortality (4, 12, 15). The microcystin content of the liver can serve as an overall indicator of fish health to better understand the consequences of HABs exposure. Measuring microcystins in both tissues using the more accurate MMPB Lemieux Oxidation method followed by liquid chromatography coupled to mass spectrometric analysis (19) addresses concerns regarding human and fish health as well as analytical limitations of many previous studies. Here, we hypothesize that the microcystin concentration of fish muscles will pose low human health risks and fish will accumulate microcystins in livers commensurate with recent HAB exposure.

## Results and Discussion

### Muscle concentration and human health risk

Fish muscles contained microcystins (F_1,184_=145.58, *p* <0.0001), at a mean concentration of 1.80 ng g^-1^ ww across species and locations (Fig. 1). However, only 2/190 of samples had concentrations exceeding the WHO’s TDI. The margin of exposure and the hazard quotient, used in human health risk assessment to estimate the level at which a compound is carcinogenic and can cause adverse effects, both suggest microcystins present a low risk (Dataset S1). Applying these results against the draft fish consumption advisory benchmarks for Lake Erie, 89% of muscles were <6 ng g^-1^ ww microcystins, the lowest fish consumption advisory benchmark in Ontario (Table S1). All muscles are in the ‘unrestricted’ meal category (<25 ng g^-1^ ww) in Ohio (Table S2). Overall, the contents of microcystins we observed in the Lake Erie fish included in our study poses no additional human health risk.

**Fig. 1.**
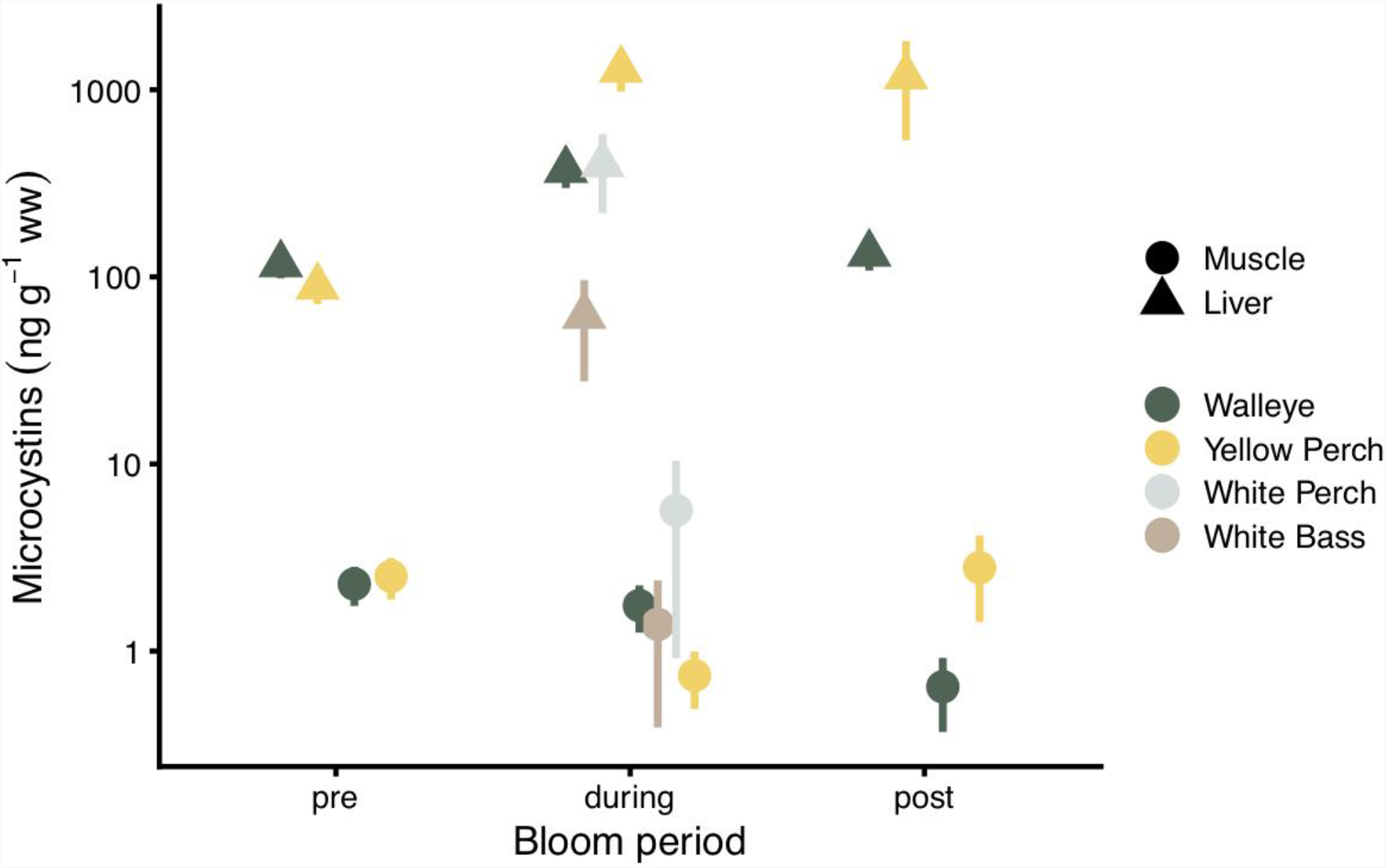
Total microcystins (ng g^-1^ ww) were measured in muscles and livers from four fish species (Walleye, Yellow Perch, White Perch, White Bass) collected pre-, during, and post-harmful algal bloom in Lake Erie.

### Liver concentration and fish health impacts

Fish accumulate microcystins in their livers (F_1,184_=184.74, *p* <0.0001), with a mean of 460.13 ng g^-1^ ww across species and locations (Fig. 1). Liver microcystin concentrations varied across HAB periods (F_2,182_=14.43, *p* <0.0001), with fish collected during the bloom having >6× the concentration of those collected pre-bloom. However, pre-bloom concentrations were still appreciable, demonstrating that fish retain toxins as a legacy of prior exposure. Fish species also differed in their liver microcystin content (F_3,183_=11.51, *p* <0.0001), with the highest concentrations in Yellow Perch. Liver concentrations of microcystins varied by location within Lake Erie (F_2,182_=39.31, *p* <0.0001), with fish caught in the western basin having 44% higher microcystins than fish from the east central basin.

The consistently high microcystin concentrations in fish livers indicate a potential biological issue not currently considered by management agencies. Acute exposure to microcystins has long been postulated to be among several stress factors involved in fish kills during HAB events (20). It is well-documented that microcystins are potent inhibitors of protein phosphatases in hepatocytes (4, 8), causing tumor-promotion, liver damage, and protein dysregulation in juvenile and adult fish even at sublethal levels (12, 21). Microcystin levels observed here are comparable with concentrations previously shown to have toxic effects during the development of embryos and larvae across a range of fish species (12). These sublethal effects suggest the concentrations we observed in wild fish may translate to negative impacts on population dynamics. Fish livers contained comparatively high values of microcystins even prior to the yearly bloom period, suggesting microcystins are accumulated and retained. Sustained microcystin levels, with peaks corresponding with seasonal HABs, reinforces the supposition that sublethal impacts on fish could have considerable effects on fish growth and recruitment. These effects could be an undocumented cause of reduced Yellow Perch recruitment (22), a major economic and management issue in Lake Erie. Planned phosphorus load reduction is predicted to increase algal toxins in Lake Erie (18) which may exacerbate any fish recruitment issues.

Concerns about toxicant exposure via human consumption are centered on muscle concentrations based on the convention of consuming fillets. Whole fish consumption is practiced in various communities due, in part, to added nutritional benefits (23). Microcystins are stable during cooking and boiling (24) and can potentially leach out of organs and spread onto fillets. These data suggest that general and sensitive populations should avoid eating whole fish from Lake Erie and other waterbodies with recurring seasonal blooms, restricting consumption to fillets.

This study demonstrates microcystins are retained in fish livers well after a HAB event and may be a persistent contaminant in aquatic ecosystems. Many fish species are highly mobile and some go on large-scale seasonal migrations within the Great Lakes, owing to factors such as foraging opportunities, behavioral thermoregulation to avoid warm waters, competitive pressure, selective fishing pressure, genetic predisposition, and reduced habitat quality due to HABs (25). Thus, the possibility exists that fish tissue concentrations may not necessarily correlate with capture location and associated water toxins, raising the profile of HABs as a wildlife and resource management issue. Our study shows that eating fish fillets from Lake Erie poses little human health risk. However, recurring and increasingly severe toxic HABs likely have major effects on freshwater fish and fisheries and deserve future study to protect the health of these ecologically and economically important species.

## Materials and Methods

Fish capture times and locations were determined following the Lake Erie Harmful Algal Bloom Forecast published biweekly by the National Oceanic and Atmospheric Administration (https://coastalscience.noaa.gov/research/stressor-impacts-mitigation/hab-forecasts/lake-erie/). Muscles and livers were analyzed for total microcystins (free and bound) using an optimized Lemieux Oxidation method (19) on individuals and, when many individuals of a given species were captured within a given timeframe, pools of individuals. A human health risk assessment was performed using microcystin concentrations for fish muscles, and the data generated was compared against fish consumption advisory benchmarks for microcystins in Lake Erie. The margin of exposure and hazard quotient were additionally calculated to estimate the potential for adverse effects to develop in humans from exposure to microcystins. Linear mixed effects (LME) models were used to test for significant concentrations of microcystins and to identify whether species identity, location within Lake Erie, and HAB status influenced concentrations. See *SI Appendix* for further details.

## Acknowledgements

This work was supported by an NSERC CREATE Grant (2013-432269), a Banting Postdoctoral Fellowship, a Canada-Ontario Agreement (2218) through the Ontario Ministry of the Environment, Conservation and Parks, and a Government of Ontario Grant (GLS 1403). Financial support by the Government of Ontario does not equal endorsement of this paper. We thank the field technicians from the Ontario Ministry of Northern Development, Mining, Natural Resources and Forestry for collecting fish from Lake Erie for this study.

## Supporting Information

Materials and Methods, Figures, Tables, References

