## Supplementary material for "Lake Erie fish safe to eat yet afflicted by algal hepatotoxins": SI Appendix

**Number of pages: 9**
**Number of tables: 2**
**Number of figures: 3**

Table of Contents


| Table of Contents | Page |
|---|---|
| SI 1. Fish capture times and locations | S3 |
| SI 2. Fish tissue analysis | S3-4 |
| SI 3. Human health risk assessment | S4-5 |
| SI 4. Statistical analysis | S5 |
| SI 5. Dataset S1 | S5 |
| Table S1. Fish consumption advisory benchmarks for microcystin-LR for general and sensitive populations in Ontario, Canada | S5 |
| Table S2. Fish consumption advisory benchmarks for total microcystins for all populations in Ohio, United States | S5 |
| Figure S1. Lake Erie Commercial Fishing Grid used for collecting fish during study | S6 |
| Figure S2. Relationship between microcystins in liver and total length of fish | S6 |
| Figure S3. Relationship between microcystins in liver and round weight of fish | S6 |
| Supplementary References | S7 |

### *SI 1. Fish capture times and locations*

A total of 190 fish from Walleye, White Bass, White Perch, and Yellow Perch were collected in 2015 and 2017 from Lake Erie (Figure S1) as a part of the Fish Contaminant Monitoring Program of the Ministry of the Environment, Conservation and Parks (Etobicoke, Ontario, Canada) and through existing fisheries assessment programs on the Lake Erie Management Unit of the Ontario Ministry of Northern Development, Mines, Natural Resources and Forestry (Wheatley, Ontario, Canada). Randomized sub-sampling from the existing fisheries assessment programs targeted fish in the following HABs periods: 'pre' (June to mid-July), 'during' (mid-July to October), and 'post' (October to November) in the western, west central, and east central basins of Lake Erie. At time of sampling, National Oceanic and Atmospheric Administration's Lake Erie Harmful Algal Bloom Bulletin was used to define the above targeted time periods. These HAB areas were based on available chlorophyll *a* and microcystin toxin concentrations. Archival records of these bulletins can be accessed here: https://coastalscience.noaa.gov/research/stressor-impacts-mitigation/hab-forecasts/lake-erie/.

Gillnetting and trawling protocols from the existing fisheries assessments do not permit precise capture locations of individual fish, thus catch locations for all collected fish were reported based on a 5-minute grid system used for Lake Erie fisheries management. Grid locations were then converted into a single GPS location based on a grid centroid. The Lake Erie Harmful Algal Bloom Bulletin was also used to classify catch locations relative to the HAB event (i.e., 'out', 'near', 'in', or 'in-peak'). Fish classified as "out" came from a 5-minute grid system using a minimum of two grids away from one containing a bloom based on the Lake Erie Harmful Algal Bloom Bulletin closest to the date of capture; "near" represents a grid where the bloom was indicated in an adjourning grid; "in" represents a grid containing part of the bloom; and "in-peak" represents a grid with high concentrations of cyanobacterial cells of the bloom.

### *SI 2. Fish tissue analysis*

As fish were collected from Lake Erie, muscles and livers were dissected and quickly frozen at -80 °C until microcystin tissue analysis. Fish tissues were analyzed for total microcystins (free and bound) using an optimized MMPB Lemieux Oxidation method that provides the necessary sensitivity and specificity (Anaraki *et al.*, 2020). Briefly, frozen tissues were freeze-dried using a Labconco Freezone 2.5 freeze-drier from Fisher Scientific (Markham, Ontario, Canada) for at least 24 h and then ground to a fine powder using a pestle and mortar. Next, approximately 100 mg of freeze-dried sample was spiked with the internal standard ($d_3$-MMPB), oxidized at pH 8.5 using potassium permanganate (0.3 mM) and sodium periodate (20 mM) for 2 h. The oxidizing sample was quenched by adjusting the pH to 3 with 10% sulfuric acid and sodium bisulphite. Samples were then centrifuged for 7 min at 5,000 × g and correspondingly loaded onto Oasis HLB 3 cc (400 mg) LP extraction cartridges (Mississauga, Ontario, Canada) for clean-up and extraction of MMPB and $d_3$-MMPB with 0.1% acetic acid and 50% methanol washes. Analytes were eluted with 4 mL of 100% methanol, dried down to 250 µL, and diluted to 1 mL using 0.1% FA milli-Q water. Samples were finally filtered using Pall GHP filters (Mississauga, Ontario, Canada) and prepared for instrumental analysis.

Samples were quantified using an *in-situ* generated MMPB matrix-matched calibration curve by isotope dilution with $d_3$-MMPB by liquid chromatography coupled to time-of-flight mass spectrometry (LC-QTOF MS) using an Acquity I Class Chromatograph by Waters Corporation (Milford, Massachusetts, United States). This approach estimates potential matrix effects that may have impacted the derivatization, sample preparation and instrumental analysis steps. This method showed 16.7% precision (RSD) and +6.7% accuracy (bias), with a calculated method detection limit (MDL) of 2.18 ng $g^{-1}$ ww.

### *SI 3. Human health risk assessment*

Total microcystins measured in muscles were compared against Ontario's fish consumption advisory benchmarks for microcystin-LR developed by the Ontario Ministry of the Environment, Conservation and Parks (Table S1) and Ohio's fish consumption advisory benchmarks for total microcystins developed by the Ohio Environmental Protection Agency (Table S2). These benchmarks are based on the World Health Organization's recommended tolerable daily intake (TDI) of 0.04 µg $kg^{-1}$ bodyweight per day (WHO, 2020). The TDI value is based on the no-observed-adverse-effect-level (NOAEL) liver pathology observed in a 13-week study on mice using microcystin-LR (Fawell *et al.*, 1994) and applying an uncertainty factor (UF) of 1000 (Eq. 1).

$$(1) \quad \mathrm{TDI} = \frac{\mathrm{NOAEL}}{\mathrm{UF}} = \frac{40\ \mu\mathrm{g\ kg^{-1} d^{-1}}}{10\ \times\ 10 \times 10} = 0.04\ \mu\mathrm{g\ kg^{-1} d^{-1}}$$

Ontario's advisory benchmark assumes that 50% of microcystin exposure is through fish consumption. Microcystin-LR is one of the most toxic microcystin congeners (Shimizu *et al.*, 2014). Therefore, application of a microcystin-LR advisory benchmark to total microcystins presents a more conservative and health protective scenario.
We further calculated the estimated daily intake (EDI) for total microcystins if one meal of fish is consumed every day (Eq. 2) and compared this to the WHO's TDI for microcystin-LR.

$$(2) \quad \mathrm{EDI} = \frac{C_{\mathrm{microcystins}} \times D_{\mathrm{intake}}}{\mathrm{bw}}$$

Where $C_{microcystins}$ is the mean total concentration of microcystins in fish tissue wet weight (Dataset S1), $D_{intake}$ is the daily fish consumption for Ontarians (one meal is 227 g) following the Ontario's Guide to Eating Ontario Fish (MECP, 2020), and bw is the body weight of an average healthy adult (assumed 70 kg). We then calculated the margin of exposure (MOE) for microcystins (Eq. 3), which is commonly used in human health risk assessment to assess for potentially genotoxic or carcinogenic compounds.

$$(3) \quad \mathrm{MOE} = \frac{\mathrm{NOAEL}}{\mathrm{EDI}}$$

Since the WHO has shown evidence that microcystins are potentially carcinogenic to humans and animals, an MOE ≥ 10,000 was considered for this study to be protective for genotoxic and carcinogenic compounds. The value of 10,000 is derived by the multiplication of two UFs; a 100-fold difference between the calculated reference point and human exposure is applied for species differences and human variability, and an additional 100-fold difference is applied for inter-individual variability (EFSA, 2005). Although it is widely documented that MOEs of 10,000 are used as a protective value in the evaluation of genotoxic and carcinogenic compounds present in food where a single toxicant is being evaluated (EFSA, 2005), the extent to which microcystins is a risk remains unclear for MOEs ≤10,000. We therefore adopted the following categories to assist with MOE classification: 1–1,000 (high risk), 1,000–10,000 (medium risk), 10,000–100,000 (low risk) (Cunningham *et al.*, 2011).

We finally calculated the hazard quotient (HQ) for microcystins (Eq. 4) as the ratio of the potential exposure to microcystins and the level at which no adverse effects are expected,

$$(4) \quad \mathrm{HQ} = \frac{\mathrm{EDI}}{\mathrm{TDI}}$$

For adverse effects from microcystin exposure, HQ ≤ 0.1 is considered negligible, HQ ≥ 0.1 ≤ 1 is considered low, HQ ≥ 1 ≤ 10 indicates some hazard, and HQ ≥ 10 indicates a high hazard.

### *SI 4. Statistical analysis*

Liver and muscle microcystin data were log transformed to improve normality. Linear mixed effects (LME) models were used to test for significant concentrations of microcystins in liver and muscle tissue, with pooled status nested within species as a random effect. LMEs were also used to identify whether species identity, location within Lake Erie, and HAB status influenced concentrations. For these LMEs the random effects used were, respectively: pooled status nested within bloom period, pooled status nested within species, and pooled status nested within species.

### *SI 5. Dataset S1*

A spreadsheet detailing microcystin tissue concentrations for the 190 fish collected from Lake Erie are provided here. For details on fish consumption advisory benchmarks, see Table S1 and S2.

**Table S1.** Fish consumption advisory benchmarks for microcystin-LR for general and sensitive populations in Ontario, Canada.

| General population | | Sensitive population [a] | |
|---|---|---|---|
| Meals per month | Microcystin-LR (ng $g^{-1}$ ww) | Meals per month | Microcystin-LR (ng $g^{-1}$ ww) |
| 0 | > 188 | 0 | > 188 |
| 1 | 94 – 188 | 0 | 94 – 188 |
| 2 | 47 – 94 | 0 | 47 – 94 |
| 4 | 23 – 47 | 4 | 23 – 47 |
| 8 | 16 – 23 | 8 | 16 – 23 |
| 12 | 12 – 16 | 12 | 12 – 16 |
| 16 | 6 – 12 | 16 | 6 – 12 |
| 32 | < 6 | 32 | < 6 |

[a] Sensitive population: Women of child-bearing age and children under 15.

**Table S2.** Fish consumption advisory benchmarks for total microcystins for all populations in Ohio, United States.

| All populations [a] | |
|---|---|
| Number of meals | Microcystins (ng $g^{-1}$ ww) |
| Do not eat | > 473 |
| 1 per month | 109 – 473 |
| 1 per week | 25 – 109 |
| Unrestricted | 0 – 25 |

[a] The following values are based on the United States Environmental Protection Agency's reference dose of 0.05 μg $kg^{-1}$ $d^{-1}$.

**Figure S1.** Lake Erie Commercial Fishing Grid used for collecting fish during study.

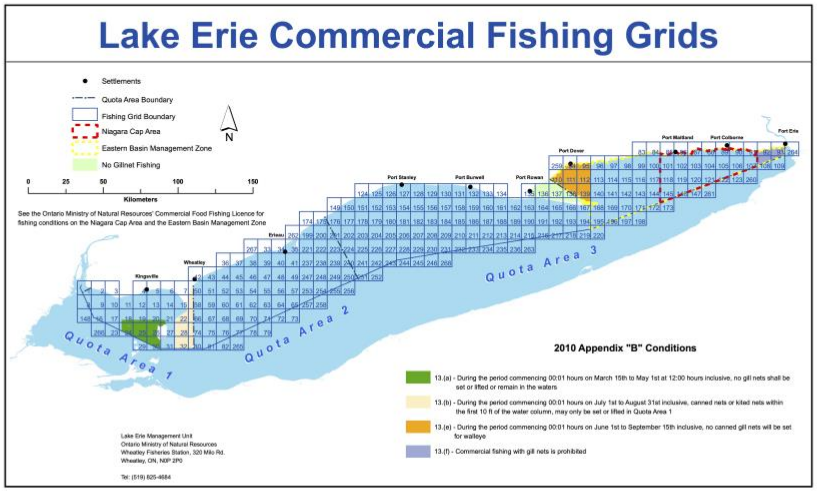


**Figure S2.** Relationship between total microcystins in liver and total length of fish.

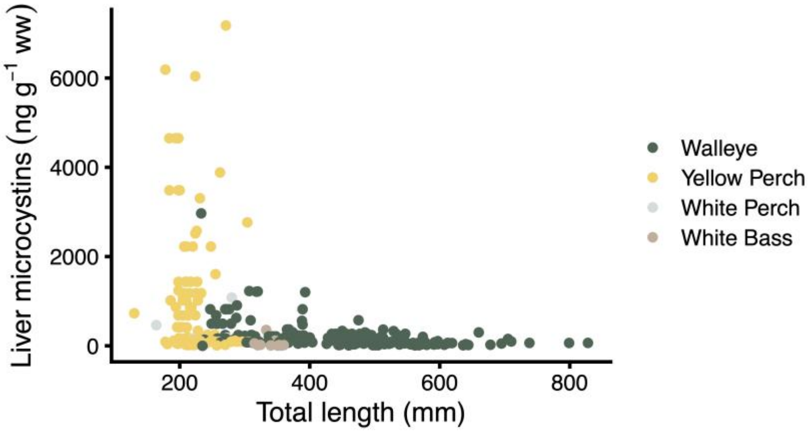


**Figure S3.** Relationship between total microcystins in liver and round weight of fish.

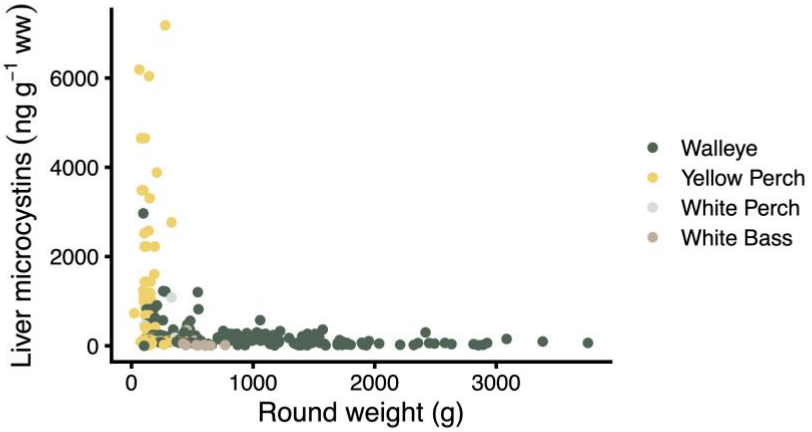
